# The marine sponge *Hymeniacidon perlevis* is a globally-distributed invasive species

**DOI:** 10.1101/2020.02.13.948000

**Authors:** Thomas L. Turner

## Abstract

In Elkhorn Slough, a tidal estuary draining into Monterey Bay, California, the intertidal is occupied by a conspicuous orange sponge known by the name *Hymeniacidon sinapium*. This same species is found in the rocky intertidal zone of the outer coast of California, and is described herein from subtidal kelp forests of Southern California. Farther afield, morphologically and ecologically indistinguishable sponges are common in estuaries and intertidal areas in Asia, Europe, South America, and Africa. Here I use morphological, ecological, and genetic data to show that these sponges are all members of the same globally-distributed species, which should be known by the senior synonym *H. perlevis*. Though previous authors have remarked upon the morphological, ecological, and/or genetic similarity of various distant populations, the true scope of this sponge’s distribution appears to be unrecognized or unacknowledged in the literature. Limited larval dispersal, historically documented range expansion, and low genetic variation all support a hypothesis that this sponge has achieved its extraordinary range via human-mediated dispersal, making it the most widely-distributed invasive sponge known to date.

**Declarations:** Conflicts of interest/Competing interests: none to declare

Availability of data and material: All raw data is included as supplementary files; georeferenced collection data is available as a supplementary .xls file; genetic data are archived at Genbank; specimen vouchers are archived at the California Academy of Sciences and at the Natural History Museum of Los Angeles; specimen photos will be made available as supplementary files, are also archived by the associated museums in GBIF, and are posted as georeferenced data on iNaturalist.org.

Code availability: n/a

## Introduction

In coastal marine ecosystems, filter-feeding marine invertebrates are among the most important invasive species in terms of species diversity, biomass, and ecological impacts (Ruiz et al. 2000; Bax et al. 2003; Byrnes and Stachowicz 2009). Sponges (phylum Porifera) are a diverse group of filter-feeding organisms that are found in all marine environments. They provide unique ecosystems services (and potential disruptions) because they preferentially consume the smaller size fractions of the plankton, such as viral and bacterial plankton (Reiswig 1971; Maldonado et al. 2012; Welsh et al. 2020). They can also have major effects on nutrient cycling, as some sponges convert dissolved nutrients into particulate matter available to other animals (de Goeij et al. 2013).

Our understanding of invasive sponges has been limited by an incomplete taxonomy. Sponges have simpler morphologies than most animals, confounding traditional classification schemes (Morrow and Cárdenas 2015). Many species were initially described as having a wide geographic range, but in recent decades these taxa have been recognized as clades comprised of multiple species with similar morphologies (Knowlton 1993; Xavier et al. 2010). This is consistent with what is known about larval dispersal in sponges. All known sponge larvae are lecithotrophic, meaning that they have no ability to feed until they settle and develop into juveniles (Maldonado 2006). They have a short planktonic stage, lasting from minutes to a few days (Maldonado 2006). Some sponges, however, do seem to have broad geographic ranges, and this is likely due to human-mediated transport. Carballo et al. (2013) list seven species thought to have recent range expansions, including two that have moved between the Pacific and Atlantic basins. Some of these species are likely to have been accidentally introduced with aquaculture (Henkel and Janussen 2011; Fuller and Hughey 2013). Trawling, hull-fouling, and other activities also likely play a role (Carballo et al. 2013).

In the current work, I describe what appears to be the most common and widely distributed invasive sponge known to date. Genetic and morphological data support a distribution that includes Europe, the Atlantic coasts of North and South America, the Pacific coast of North America, and Asia. Morphological data suggest it is also present in New Zealand, Southwest Africa, and the Pacific coast of South America, but genetic data are not yet available from these populations. In much of this range, it is among the most common sponges in multiple habitats. In Europe, this species is known as *Hymeniacidon perlevis* (Montagu 1814). The range of *H. perlevis* was already thought to be substantial: from Norway in the North to the Macronesian Islands off Africa in the South (Erpenbeck and Van Soest 2002). Within this range it is found in diverse habitats, including both the intertidal and the subtidal zones, and it can grow buried in sediment or on hard substrate (Erpenbeck and Van Soest 2002). It is often abundant in these habitats, and is considered to be one of the most common sponges in Europe (Erpenbeck and Van Soest 2002). It has been described by other taxonomists as also occurring in New Zealand (Bergquist 1961, 1970) and as the most abundant intertidal sponge in Western South Africa (Samaai and Gibbons 2005), but these records were rejected from the consensus view (Van Soest et al. 2020a), probably because limited dispersal ability seemed to make such a range implausible. Sponges from additional parts of the globe have been described as morphologically indistinguishable from *H. perlevis,* but in these cases taxonomists put forth other names for these distant populations. For example, de Laubenfels described a sponge he named *Hymeniacidon sinapium* from California in 1930 (de Laubenfels 1930, 1932). He acknowledged that “it is doubtful whether this is a new form”, and went so far as to suggest that species with the names “*sanguinea*, *luxurians*, *caruncula*, *heliophila*, *sinapium*, and perhaps even more species” are in fact synonyms. Consistent with this prediction, the European species *sanguinea* and *caruncula* have been synonymized with

*H. perlevis* (Van Soest et al. 2020a). The status of *H. luxurians* is unclear (Van Soest et al. 2020b), but the other two species, *H. sinapium* and *H. heliophila*, are still considered valid. In the current work, I will present evidence that *H. sinapium* is conspecific with *H. perlevis*, and that most sponges placed under the name *H. heliophila* are also *H. perlevis*.

When describing *H. sinapium* in California, de Laubenfels remarked on its impressive ecological breadth. He described it as abundant in the “surf-beaten” intertidal throughout Southern California, but also the most abundant sponge on the oyster beds in Newport Bay (de Laubenfels 1932). He reported only one sample from subtidal depths, but his subtidal sampling was limited, primarily via trawling. In contrast to this abundance in Southern California, de Laubenfels was only able to locate a single specimen of this species in Central California. This is notable because he was based at Hopkins Marine Station in Monterey Bay (Central California) in the 1920s, and this was the area that he studied most comprehensively at that time. A monographic report on Elkhorn Slough, which drains into Monterey Bay, was published in 1935: it reports 4 species of sponges in the estuary, but none similar to *H. sinapium* (MacGinitie 1935). This makes it unlikely that this species was present in large numbers in Central California in the 1920s.

Subsequently, however, it has become a common species in intertidal portions of Elkhorn Slough, which drains into Monterey Bay (Wasson et al. 2001), and it is also known from Tomales Bay in Northern California (Wasson et al. 2001; Fuller and Hughey 2013).

Morphological (Sim 1985) and genetic (Hoshino et al. 2008) comparisons later confirmed that a common *Hymeniacidon* in Korea, Japan, and China were the same species as those in California, so it was proposed that *H. sinapium* was introduced to California from Asia with oyster mariculture (Fuller and Hughey 2013). Though this is certainly possible, the data I compile here illustrates that it may also be non-native in Asia. This species has been said to occur in the Mexican Pacific (Hofknecht 1978) and the Galapagos Islands (Desqueyroux-Faúndez and Van Soest 1997) as well, but genetic data are not yet available from those populations.

The final species to consider, *H. heliophila* (Wilson 1911), is ascribed a substantial range in the Western Atlantic, from the Gulf of Maine to Brazil (Muricy and Hajdu 2006; Weigel and Erwin 2016; Van Soest et al. 2020c). Originally described as the most abundant sponge in Beaufort Harbor North Carolina (Wilson 1911), it is also said to be very common in the Caribbean (Diaz et al. 1993). A recent paper also found that an indistinguishable sponge was the most common intertidal sponge present in the Bahía San Antonio, Argentina, (Gastaldi et al. 2018). In this case, the authors opted to refer to their samples by the name *H. perlevis*, as the Argentinian samples were indistinguishable from ones in Northern Europe in genotype, habitat, and morphology (Gastaldi et al. 2018). Here, I build on these results by 1) analyzing additional samples from Southern California, which contains the type locality for *H. sinapium*, and 2) compiling all publicly available genetic data (from 20 publications and several unpublished datasets). When presented together, the data provide a compelling case for a single species ranging across both the Atlantic and Pacific basins and the Northern and Southern hemispheres. Given the limited dispersal capabilities of the species (Xue et al. 2009), the limited genetic variation over most of its range (see below), and the historically documented range expansion in California, these data are most consistent with an invasive spread via human facilitation.

## Methods

### Collections

Sponges were located while SCUBA diving. Effort was made to photo-document all sponge morphotypes present at each dive site, so that data on presence vs. absence could be compiled. It should be noted, however, that search time was higher at some sites than others, as shown in supplementary table 1. The search times listed are the total dive time, cumulative across all dives at a site. This only approximates search effort, as some dives were spent mainly searching and photographing sponges, while on others considerable time was spent collecting samples. Collections were made by hand with a small knife. Samples were placed individually in plastic bags while underwater, accompanied with copious seawater. These bags were put on ice until preservation, which was generally within 2-5 hours, but sometimes up to 12 hours. Samples were moved directly from seawater to 95% ethanol; in most cases, the preservative was changed with fresh 95% ethanol after 1-3 days, and sometimes changed again if it remained cloudy. Most samples were photographed underwater with an Olympus TG5 before collection and photographed again in the lab. These photos (and the microscope images discussed below) accompany this paper as supplementary data and are also posted as georeferenced records on the site iNaturalist.org. Two samples were collected during the “LA Urban Ocean Bioblitz”, and are present as vouchers at the Natural History Museum of Los Angeles. Three other samples were deposited with the California Academy of Sciences in San Francisco. Voucher numbers are shown in supplementary table 1. This table lists all samples known or suspected to be *Hymeniacidon sinapium*. Note that the standard of evidence is variable in each case, as indicated in the table (e.g. some were photographed but not collected, and the ID is therefore tentative; see results for further details).

### Spicules

Sponge spicules were examined by digesting soft tissues in bleach. A subsample of the sponge was chosen, taking care to include both the ectosome and choanosome. This was placed in a 1.5 ml microcentrifuge tube with household bleach for several hours, until tissue appeared to be dissolved. With the spicules settled at the bottom of the tube, the bleach was then pipetted off and replaced with distilled water; this was repeated several times (I found that 2-3 water changes were sufficient for visualizing spicules with a light microscope, but removing all the salt from the sample for other downstream applications required 5 or more rinses and worked best when the final ones were done with absolute ethanol). In some cases, samples were centrifuged at low speed to reduce settling time between rinses, though this increased the proportion of broken spicules.

Spicules were imaged using a compound triocular scope and pictures were taken using a D3500 SLR camera (Nikon) with a NDPL-1 microscope adaptor (Amscope). Pictures of a calibration slide were used to determine the number of pixels per mm, and 20-30 spicules were then measured using ImageJ (Schneider et al. 2012). Spicules length was determined in a straight line from tip to tip, even when spicules were curved or bent. Spicules were selected randomly, so as to get an unbiased estimate of size distributions. All raw data are available as supplementary table 2. I also imaged the spicular architecture in cleared tissue sections. I used a razor blade to cut perpendicular sections that were as thin as possible by hand, and removed the surface layer (ectosome) by hand for a surface view. These sections, already in 95% ethanol, were soaked in 100% ethanol for a short time and then cleared for several hours in Histoclear (National Diagnostics).

### Genotyping

DNA was extracted from small subsamples, taking care to minimize contamination by the many sponge-associated organisms that are often present. Some samples were extracted with the Qiagen Blood & Tissue kit while the others were extracted with the Qiagen Powersoil kit. The “barcoding” region of the cox1 gene was amplified using the Folmer primers LCO1490 (GGTCAACAAATCATAAAGAYATYGG) and HCO2198

(TAAACTTCAGGGTGACCAAARAAYCA) (Folmer et al. 1994). A single attempt was made to amplify a longer region using the primers from Rot et al. (Rot et al. 2006): LCO1490 and COX1-R1 (TGTTGRGGGAAAAARGTTAAATT) but without success. A portion of the 18S locus was amplified using the primers SP18aF (CCTGCCAGTAGTCATATGCTT) and 600R18S (CGAGCTTTTTAACTGCAA) (Redmond et al. 2013); the C2-D2 region of 28S was amplified using primers C2 (GAAAAGAACTTTGRARAGAGAGT) and D2 (TCCGTGTTTCAAGACGGG) (Chombard et al. 1998). All primer sequences are listed 5’ to 3’. PCR was performed in a Biorad T100 thermocycler with the following conditions: 95C for 3 min, followed by 35 cycles of 94C for 30 sec, 52C for 30 sec, 72C for 1 min, followed by 72C for 5 minutes. The C2-D2 28S region was amplified with a 57C annealing temperature instead of 52C. PCR was performed in 50 ul reactions using the following recipe: 24 μL nuclease-free water, 10 μL 5× PCR buffer (Gotaq flexi, Promega), 8 μL 25mM MgCl, 1 μL 10mM dNTPs (Promega), 2.5 μL of each primer at 10 μM, 0.75 μL bovine serum albumin (10 mg/ml), 0.25 μL Taq (Gotaq flexi, Promega), 1 μL template. ExoSAP-IT (Applied Biosystems) was used to clean PCRs, which were then sequenced by Functional Biosciences (Madison, Wisconsin) using Big Dye V3.1 on ABI 3730xl instruments. All PCR products were sequenced in both directions, and a consensus sequence was constructed using Codon Code v.9 (CodonCode Corporation). Blastn was used to verify that the resulting traces were of sponge origin; all sequences have been deposited in Genbank as accessions MT007958-MT007960 (cox1), MT001298 (18S), and MT006362 and MT422190 (28S). See supplementary table 3 for additional details and information.

### Genetic analysis

I retrieved all sequences with high sequence similarity to *H. perlevis* from Genbank. I used the NCBI taxonomy browser to compile all data from samples identified as *H. perlevis, H. sinapium, H. heliophila*, and *H. flavia.* Together, these data are from 20 different publications and several datasets that were deposited in Genbank but never published (Erpenbeck et al. 2002, 2004, 2005, 2006, 2007; Park et al. 2007; Hoshino et al. 2008; Turque et al. 2008; Erwin et al. 2011; Alex et al. 2012, 2013; Morrow et al. 2013; Redmond et al. 2013; Thacker et al. 2013; Fuller and Hughey 2013; Jun et al. 2015; Miralles et al. 2016; Weigel and Erwin 2016; Gastaldi et al. 2018; Regueiras et al. 2019). I also retrieved all samples identified as *Hymeniacidon* sp. and checked these and other sequences using blastn for similarity to *H. perlevis/sinapium/heliophila*. Only four of these (JN093018 and KU697715-KU697717), all identified as *Hymeniacidon* sp., were closely related to the other samples, and all appeared to be identical to other sequences within the *H. perlevis* clade. These four unidentified samples were not included in downstream analyses.

I was not able to use all sequences in every analysis because of differences in the sequenced portion of the gene or a lack of information regarding collecting location. Importantly, no samples were excluded simply because they showed discordant patterns of sequence variation. Supplementary table 3 lists every Genbank accession found, indicates which were included in each analysis, and explains the reasons why any were excluded. Some reads were included in the phylogenetic analysis, which could tolerate unequal read lengths, but not the haplotype network, which included only samples with complete data over the entire alignment. Sequence alignments were produced in Codon Code v.9 (CodonCode Corporation). Haplotype networks were produced using the minimum spanning method (Bandelt et al. 1999) as implemented in Popart (Leigh and Bryant 2015). Phylogenies were estimated with maximum likelihood using IQ-Tree (Nguyen et al. 2015; Trifinopoulos et al. 2016). I used a GTR model of sequence evolution and used the Ultrafast bootstrap (Hoang et al. 2018) to measure node confidence. Phylogenies were produced from the IQ-Tree files using the Interactive Tree of Life webserver (Letunic and Bork 2019). Figures were made ready for publication using R (r-project.org) and/or Gimp (gimp.org).

## Results

### Status in Southern California

Little data has been published about the distribution of *Hymeniacidon sinapium* in California outside of bays and estuaries. Past surveys have focused on intertidal habitat and/or subtidal sampling via deep water trawl (de Laubenfels 1932; Sim and Bakus 1986; Bakus and Green 1987; Green and Bakus 1994). It was therefore unknown if *H. sinapium* is present in kelp forest ecosystems. I searched for it using SCUBA at 47 sites in Southern and Central California (table S1). Subtidal sites were shallow rocky reefs except for two locations which were oil platforms. Subtidal sites include four marine protected areas along the mainland coast and three marine protected areas within the Channel Islands National Marine Sanctuary. Six of the sites are also field sites within the Santa Barbara Coastal Long-Term Ecological Research Network (sbclter.msi.ucsb.edu). Though the survey was focused on kelp forest habitats, I also checked two intertidal sites and floating docks in two harbors, as shown in table S1.

The distribution of *H. sinapium* in the Channel Islands region, where sampling was most comprehensive, is shown in figure 1. I found the sponge at 8 of 19 mainland reefs in Southern California, including both mainland marine protected areas investigated in Southern California (Naples and Campus Point Marine Protected Areas). In only one location (Carpinteria Reef) did I find *H. sinapium* growing on rock; in all other locations it was largely buried in sediment, with projections extending into the water column. In contrast to its prevalence on the Southern California mainland, I did not find it at any island sites. This difference seems unlikely to be due to dispersal limitation because island and mainland sites have high connectivity (Watson et al. 2010). It is more likely due to the ecological differences between sites: none of the island sites investigated had areas with the fine silty sediment where the sponge was most common on the mainland. Though silty sites at the islands may simply have been unsampled in this survey, it is likely they are less common than on the mainland. For example, satellite data show that particles at the islands are less prone to resuspension by wave action (Freitas et al. 2017). An intertidal survey of island sites in the 1970s did find *H. sinapium* at both San Miguel and Santa Rosa Islands (Bakus and Green 1987). It has also been reported from the more Southern islands of Catalina and San Clemente, which were barely sampled by my survey (Sim and Bakus 1986; Bakus and Green 1987). To the North, in Central California, I only surveyed three subtidal sites. I did not find *H. sinapium* in any of these Central California sites, nor did I find it at the few intertidal sites, floating docks, or oil rigs that were checked.

**Figure 1.**
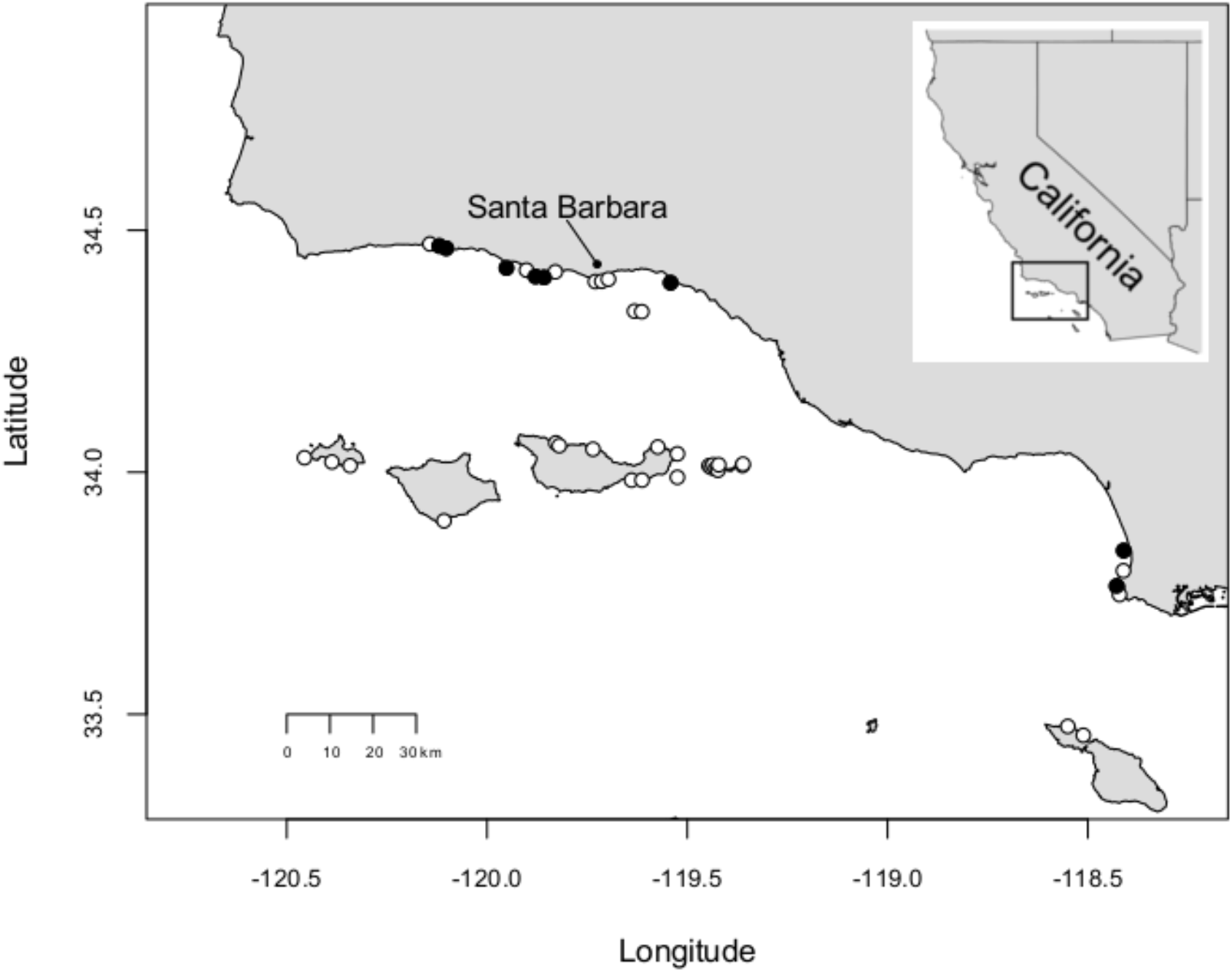
Collection locations in the Southern California Channel Islands region. Sites where *H. sinapium* were found (black) and not found (white) are shown. The two sites away from the coastline are oil platforms. Collection sites in Central California are not shown.

Together, my recent collections and the published intertidal and bay surveys in California produce a portrait of a species that thrives in a wide variety of conditions, from bays to the rocky intertidal to the kelp forest (Lee et al. 2007). It seems most abundant in the intertidal in some bay habitats with a muddy substrate and high sedimentation, and seems more common in the kelp forest where fine sediment is found. These data are completely consistent with the published descriptions of habitat preferences for *H. perlevis* in Europe (Erpenbeck and Van Soest 2002) and *H. heliophila* in the Western Atlantic (Weigel and Erwin 2016).

### Gross Morphology of kelp forest samples

All but one of the newly collected samples were found embedded in sediment with irregular projections extending into the water column. These projections varied from stout cone-shaped or bag-shaped oscula to long, tendril-like digitations. One sponge was found unburied, growing on rock. It lacked projections and instead resembled *Halichondria panicea* (its identity was confirmed with spicule and DNA data, presented below). All samples had a fleshy consistency, with the rock-dwelling sponge somewhat firmer. Color varied from yellow to yellowish-orange in the field. Field photos are available for 8 samples in the supplementary data accompanying this paper, and are also available at iNaturalist.org.

I was interested in whether these sponges could be identified in the field and therefore monitored using roving diver surveys or photo transects. These samples were collected as part of an ongoing project to characterize the diversity of kelp forest sponges, with over 500 samples collected to date. This is one of the first surveys of sponges in California via SCUBA, and the first with extensive field photos of specimens that have also been analyzed morphologically. Though the bulk of these data will be published elsewhere, comparisons to date indicate that *H. sinapium* is the only sponge in these habitats that grows by extending irregularly shaped projections out of silty sediment. Though this morphology is certainly known from other species, *H. sinapium* was the only sponge with this morphology found within the sampling effort shown in supplementary table 1. This indicates that this morphology, when found in the Southern California kelp forest, is strongly suggestive of the presence of this species. The most similar species found to date is *Polymastia pachymastia*: as the name suggests, this sponge is covered in nipple-like projections. This sponge was also found covered in sediment, with only the projections visible. However, these projections tend to be uniform in shape and regularly spaced in *P. pachymastia*, which contrasts with the irregularly spaced and morphologically various projections seen in *H. sinapium.* The projections are also nearly white in *P. pachymastia*, while they vary from yellow to nearly orange in *H. sinapium.* The rock-dwelling *H. sinapium* found at Carpenteria Reef, however, would be more challenging to identify from field photos, as it is very similar to other Halichondridae found in the survey.

### Spicular morphology

I characterized the spicules of 9 samples to confirm their identity and compare them to published data. All spicules were styles: tapered to a point at one end, and rounded at the other. Width was usually uniform over the entire length, but a small minority had faint swelling at or near the rounded end. This was manifest as a very weak swollen head including the end (similar to the head of a match), or more commonly as a swollen band near the head end (like a bead on a string). Most were somewhat curved or bent. The skeleton of one sample was investigated further using hand-cut sections cleared with Histoclear. Spicules in perpendicular sections through the choanosome formed wavy, meandering tracts, the largest of which were about 30 μm wide. Spicules were also found outside the tracts pointing in all directions (referred to as a “confused” arrangement in sponge taxonomy). Surface sections revealed that the ectosome of the sponge was filled with spicules that appeared to be tangential (parallel to the sponge surface) and also “paratangential” (at an angle to the surface of less than 90 degrees). These spicules were in messy bundles that formed an approximate mesh on the surface of the sponge. Table 1 shows measurements of spicules as compared to values published in other studies of *Hymeniacidon*. Newly collected data are consistent with published data from *H. sinapium, H. perlevis,* and *H. heliophila,* as well as *H. fernandezi* (Thiele 1905) from Chile (for which no genetic data is yet available). The arrangement of spicules in cleared sections is also congruent with the spicular architecture described for *H. perlevis* and other species (Erpenbeck and Van Soest 2002). Photos of tissue sections and spicules are available as supplementary data.

**Table 1.** Morphological data from newly collected samples with comparison data from the literature.

| Sample | Date | Collection location | N | Style length | Style width |
| --- | --- | --- | --- | --- | --- |
| TLT550 | 1/7/20 | Arroyo Quemado | 25 | 151 - 262 - 336 | 3 - 5 - 7 |
| TLT79 | 7/1/19 | Tajigus | 26 | 121 - 223 - 322 | 3 - 5 - 7 |
| TLT87 | 7/1/19 | Tajigus | 21 | 147 - 216 - 326 | 2 - 5 - 8 |
| TLT109 | 7/1/19 | Tajigus | 32 | 138 - 289 - 415 | 4 - 6 - 8 |
| TLT339 | 8/30/19 | Coal Oil Point | 25 | 160 - 283 - 376 | 2 - 5 - 6 |
| TLT247 | 7/31/19 | Carpinteria | 20 | 129 - 223 - 319 | 4 - 5 - 7 |
| TLT129 | 7/31/19 | Carpinteria | 36 | 116 - 246 - 334 | 3 - 5 - 7 |
| TLT15955 | 8/22/19 | Redondo Barge | 25 | 127 - 235 - 407 | 3 - 5 - 7 |
| TLT349 | 8/23/19 | Resort Wall | 23 | 163 - 264 - 351 | 2 - 5 - 12 |
| *H. sinapium*<sup>1</sup> | - | Elkhorn Slough, CA | - | 115 - 460 | 3 - 12 |
| *H. perlevis*<sup>2</sup> | - | Wales | - | 152 - 475 | 3 - 12 |
| *H. perlevis*<sup>3</sup> | - | South Africa | - | 155 - 337 | 7 |
| *H. perlevis*<sup>4</sup> | - | New Zealand | - | 189 - 329 | 2 - 10 |
| *H. heliophila*<sup>5</sup> | - | Caribbean | - | 130 - 450 | 3 - 10 |
| *H. fernandezi*<sup>6</sup> | - | Chile | - | 200 - 340 | 3 - 10 |

Date = collection date, N = number of spicules measured, spicule dimensions given in format min - mean - max; when only two numbers are shown they are min - max. Sources: 1: Lee et al. 2007; 2: Erpenback & Van Soest 2002; 3: Samaai & Gibbons 2005; 4: Bergquist 1970; 6: Thiele 1905.

### Genetic analysis

I sequenced the newly collected samples at the cox1 locus (3 samples), the 18S rDNA (1 sample) and the 28S rDNA (2 samples). A sample of the *Hymeniacidon sinapium* holotype was also loaned to me by the Smithsonian Natural History Museum: despite repeated attempts, I was unable to amplify DNA from this sample. This was not surprising, as it was collected in 1926 and little is known about its initial preservation.

I mined Genbank for all DNA data available for *H. perlevis, H. sinapium,* and *H. heliophila* (see methods). I generated sequence alignments for four loci: cox1, 18S rDNA, 28S rDNA, and a locus spanning the first intergenic transcribed spacer, the 5.8S rRNA, and the second intergenic transcribed spacer (hereafter referred to as the ITS). No other locus had more than 2 sequences available in Genbank from any of these taxa. Preliminary phylogenies indicated that sequences of *Hymeniacidon flavia* were more closely related to the clade containing my target species than anything else in Genbank. When available, these sequences were included for comparison.

Figure 2 shows the haplotype networks for the three loci with the most data. A large dataset was available for 226 base pairs at the ITS locus. A set of 271 sponges from Japan and Korea contained little genetic variation, as previously described (Park et al. 2007; Hoshino et al. 2008). Samples from Northern, Central, and Southern California were all identical to the most common Asian haplotype, as were 9 of 10 samples of *H. heliophila* from the Eastern United States. These include samples from Alabama, Florida, and North Carolina. This last sequence read is the only one available that is identified as coming from this state, which contains the type location for *H. heliophila*. As a useful comparison to the diversity in 298 samples of *perlevis/heliophila/sinapium*, a population sample of 212 *H. flavia* are shown. These are all from Japan, yet they contain a similar amount of diversity as the worldwide sample from *H. perlevis/heliophila/sinapium*. In contrast to the lack of divergence between *H. perlevis, H. heliophila,* and *H. sinapium,* the *H. perlevis/heliophila/sinapium* clade is well demarcated compared to other species. Sequences from closely related *H. flavia* differ by 8.4—9.7%. Attempts to align other published Halichondridae sequences at this locus failed due to very high sequence divergence.

**Figure 2.**
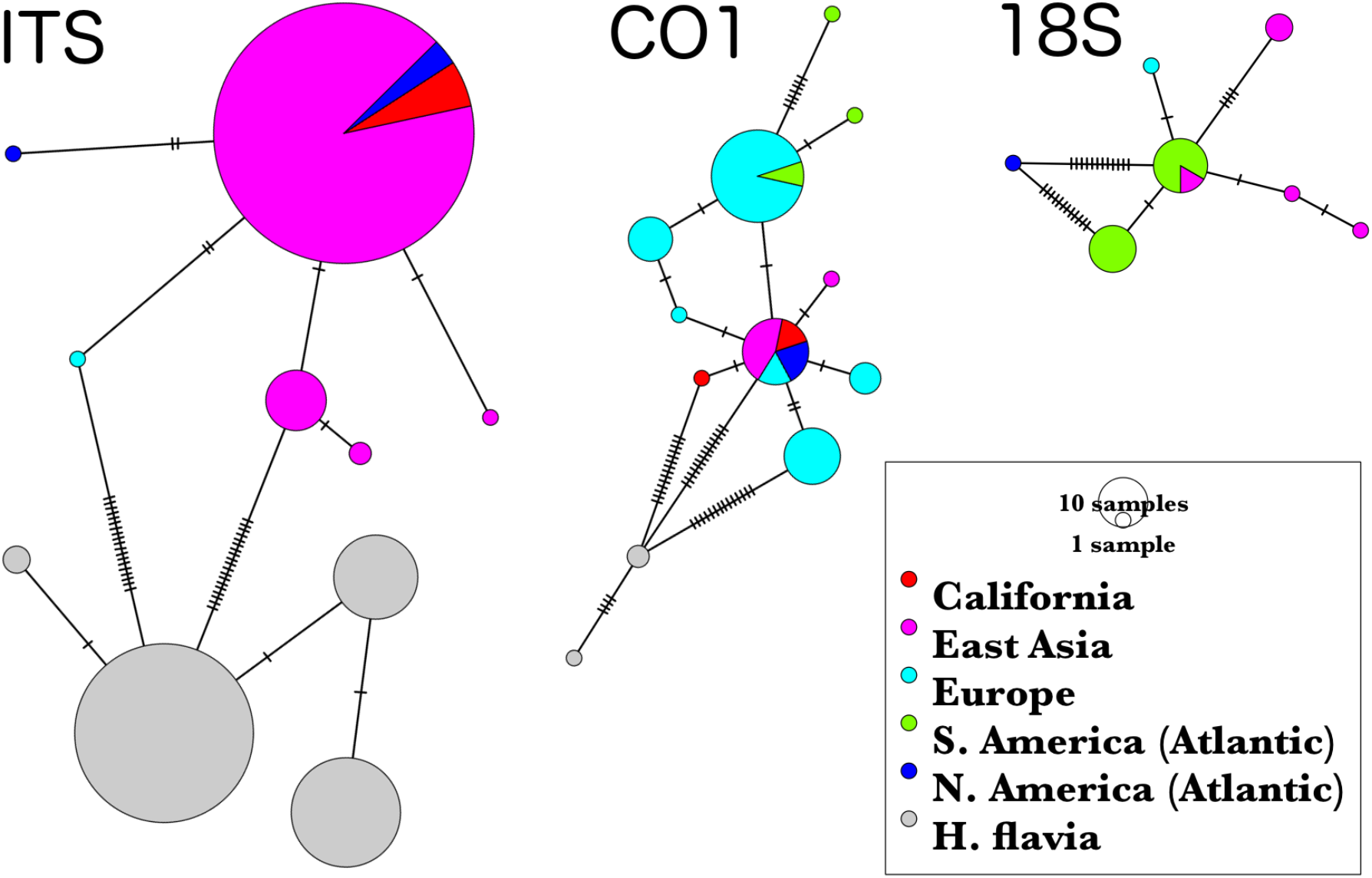
Minimum-spanning genotype networks for three loci. Samples are coded by collection location, regardless of whether they were identified as *H. perlevis, H. sinapium,* or *H. heliophila*. Closely related *H. flavia* are shown for comparison where available; all data for this species is from Japan and Korea.

A large mitochondrial dataset is also available at the Folmer barcoding region of the cox1 locus (571 bp; fig. 2). A single haplotype was shared among populations from China, Korea, Southern California, Florida, and Portugal. Samples from Argentina contained only 1–2 differences (99.6% identity) compared to this world-wide haplotype. The only sample that is more than 0.5% divergent from this common haplotype, out of all *H. perlevis/sinapium/heliophila* available, is a single sequence from Rio de Janeiro, Brazil (1.2% divergent; top of fig. 2). No morphological description of this sample is available in the related publication (Turque et al. 2008), but it states that vouchers were deposited in the Museu Nacional, Universidade Federal do Rio de Janeiro. Personal communications with Guilherme Muricy at the Museu Nacional indicate that this sample matches *H. heliophila* in both gross morphology and spicular morphology, as described in Muricy and Hajdu 2006 (p. 53). This sample is discussed further below.

Though I found that *H. perlevis, H. heliophila, and H. sinapium* shared an identical genotype at 571 bp of cox1, the cox1 locus is known to have a slower evolutionary rate in sponges than most other animals (Huang et al. 2008). Additional interspecific sequence comparisons provide context for this lack of divergence. First, the *H. perlevis/heliophila/sinapium* clade is differentiated from its closest relative, *H. flavia*, with 2.5 - 4.5% sequence divergence (2.5 - 3.7% if the sample from Rio de Janeiro is excluded). Additionally, Genbank data illustrate that most named species in the Suberitida are genetically distinguishable at cox1. Excluding the *H. perlevis/heliophila/sinapium* clade, a cox1 sequence is available from 58 vouchers across 39 species for the order Suberitida. I determined sequence divergence between each voucher and the most similar sequence from a different species. In only one case are two sequences identical over a continuous 500 bp or more. This case is for sequences identified as *Suberites pagurorum* and *Suberites domuncula*, which are themselves members of a species complex in need of taxonomic revision (Solé-Cava and Thorpe 1986). Across all 58 comparisons, average sequence divergence to the most similar conspecific voucher was 3.7% (standard deviation = 3.2%).

Less data is available for the 18S and 28S rDNA loci, but the 18S locus once again illustrates the genetic similarity of Atlantic *H. perlevis/heliophila* populations and Pacific *H. sinapium* populations (figure 2). Over the aligned 1,642 bp, samples from China shared an identical haplotype with samples from Argentina. A sample of *H. perlevis* from Ireland differs from this common haplotype by only a single base pair (this is the closest available data to the type locality for *H. perlevis*, which is the Devon Coast in England). Only a single data point has any notable divergence: a sponge identified as *H. heliophila* from the USA. This sample is separated from all others by 12 substitutions (0.7% divergence). I created a phylogeny including selected Halichondridae to place this divergence in context (figure 3). While all other sequences of *H. perlevis/heliophila/sinapium* form a clade, this USA sample is as divergent as other distinct species. The interior nodes of this phylogeny are not well resolved, but it is clear that this sequence is an outlier and likely from a different species. This sample (NCI217, Smithsonian voucher #0M9G1074-H) is part of a collection of sponges for the National Cancer Institute deposited at the Smithsonian Museum of Natural History (Redmond et al. 2013). It was collected by the Coral Reef Research Foundation in the Florida Keys (Key Largo, from mud substrate), and identified by Belinda Glasby (William Moser, Smithsonian Museum of Natural History, pers. comm.). It is discussed further below. Divergence between all other sequences in the *H. perlevis/heliophila/sinapium* clade and the most closely related species available (*Halichondria bowerbanki*) is 1.4 - 1.8%. This is less divergence than at the ITS and cox1 loci, indicating lower power to discriminate among species on a per-base basis. The aligned region is 3 times longer than cox1 and 7 times longer than ITS, however.

**Figure 3.**
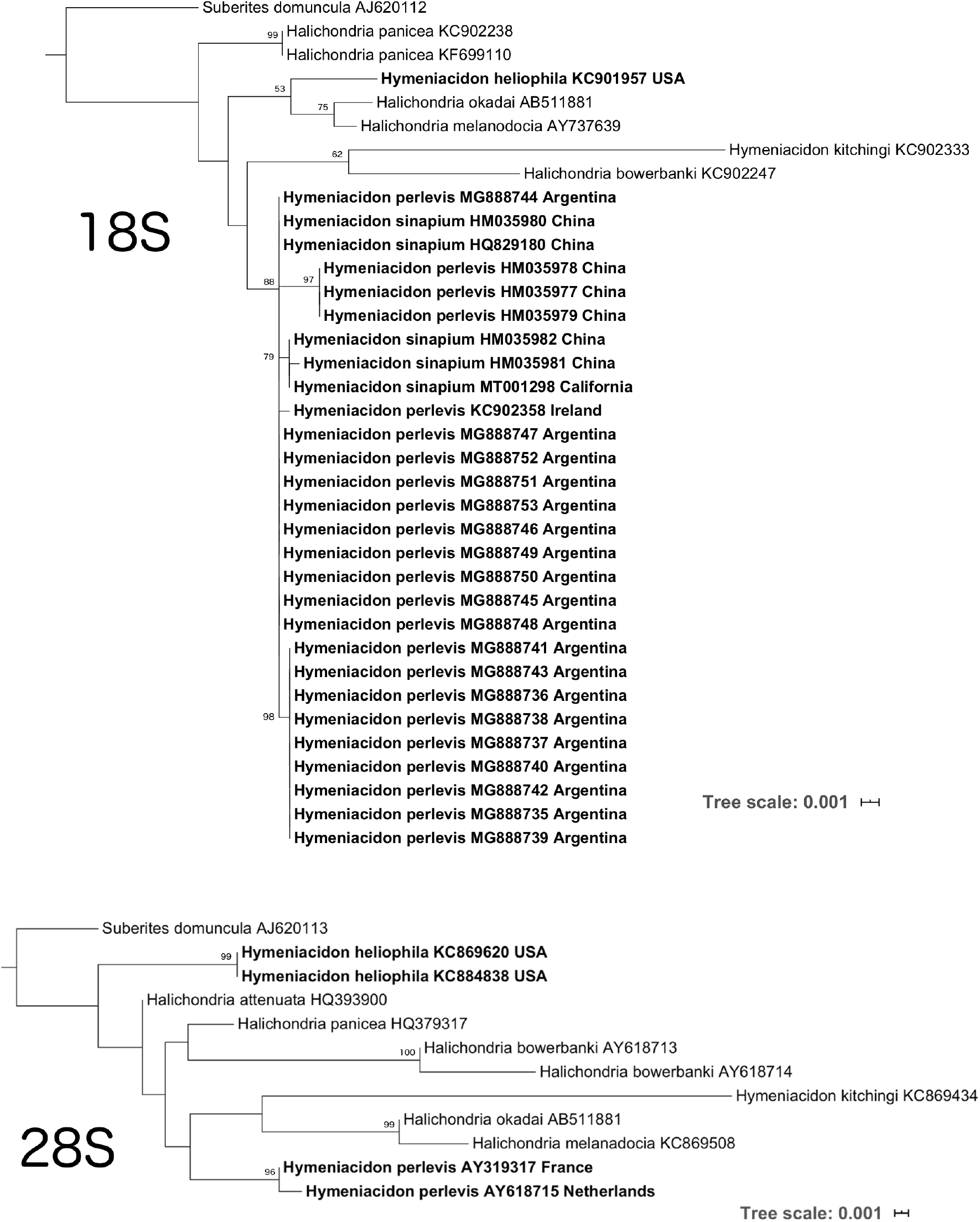
Phylogenies for the 18S and 28S loci. All samples identified as *H. perlevis, H. sinapium* and *H. heliophila* are shown in bold with localities. Selected other Halichondridae are shown for comparison, with *Suberites domuncula* specified as the outgroup. Genbank accession numbers are also shown. Ultrafast Bootstrap support is shown for all nodes with > 50% support.

The D3–D5 region of 28S also allowed for an interesting comparison (figure 3). The only data available at this locus is from two European samples and two from the Florida Keys, USA. One of the Florida sequences is from the same isolate as the outlier at 18S (NCI217), while the other sample is from the same collection (NCI083, Smithsonian voucher #0M9G1369-A) (Thacker et al. 2013). It was collected in the Florida Keys (Marquesas Key, sand substrate; William Moser, pers. comm.) In agreement with the 18S data, these samples do not appear to be from the same species as the European ones.

## Discussion

Genetic data provide strong support for the synonymy of *H. perlevis* and *H. sinapium*. The type locality for *H. sinapium* is Newport Bay, in Southern California (de Laubenfels 1932). Previously, the only genetic data from Southern California was from a single sample from Mission Bay, roughly 140 km to the South of the type location. To this I have added additional data from Santa Barbara County (200 km North of the type location). All of these samples are genetically identical to samples of *H. perlevis*. Indeed, there is no appreciable genetic divergence between any sample from California, Japan, Korea or Europe. One of the loci investigated, the ITS locus, is the fastest-evolving locus regularly used in sponge systematics. It evolves too fast to be informative above the level of family (Wörheide et al. 2004), and is more commonly used to infer population structure within species (Wörheide et al. 2002, Duran et al. 2004). The cox1 locus evolves more slowly, but I found that it still shows an average of 3.7% sequence divergence within this order of sponges, and a published data spanning phylum Porifera found an average of 4.9% divergence (Huang et al. 2008). The 18S locus evolves more slowly still, but the longer alignment at this locus was sufficient to differentiate other species in the family, as shown in the phylogeny. It remains possible that genomic analyses could reveal reproductively isolated groups that are not differentiated at these particular genes (e.g. Turner et al. 2008). However, there is no reason to assume any such “cryptic species” would be associated with the geographic regions previously ascribed to *H. perlevis* and *H. sinapium*. I therefore formally propose that *H. sinapium* de Laubenfels (1930) be considered a junior synonym of *H. perlevis* Montagu (1814).

It is possible that *H. heliophila* is also a junior synonym of *H. perlevis,* but some ambiguity remains. Genetic data illustrate that the majority of samples identified as *H. heliophila* are in fact *H. perlevis*, including the only one from North Carolina, which contains the type location. The two National Cancer Institute vouchers from Florida, however, appear to be from a different species. One cox1 sequence from Brazil is also modestly divergent, and could be from another species. It is possible that there are two morphologically similar *Hymeniacidon* within the range ascribed to *H. heliophila*, mirroring the case of *H. sinapium* and *H. flavia* in Japan and Korea. Though further work will be required to determine if the name *H. heliophila* is valid, is it clear that the most common sponge matching its description is in fact *H. perlevis*, whose range therefore includes North Carolina, Florida, Alabama and Argentina at the very least.

Ecological and morphological data also support these far-flung populations being within the same species. It should be noted, however, that there is another species that is morphologically similar yet genetically distinct. The genetic outgroup to the *H. perlevis/heliophila/sinapium* clade is *H. flavia*, known from Japan and Korea (Park et al. 2007; Hoshino et al. 2008). This species is sympatric with *H. sinapium* in Japan and Korea, and cannot be distinguished from it based on spicular morphology. These species can only be identified using genetic data, color when alive, or larval morphology (Sim and Lee 2003; Hoshino et al. 2008). This illustrates that it may be difficult to resolve the taxonomy of *Hymeniacidon* without genetic data. As pointed out by Gastaldi et al. (Gastaldi et al. 2018), there are additional species with morphological descriptions matching *H. perlevis*, but the existence of *H. flavia* shows that confidently determining which are synonyms will require DNA data.

Data on larval biology are available for *H. perlevis*, and they support a hypothesis of recent range expansion via human facilitation. Larvae of *H. perlevis* are non-tufted parenchymella, and do not appear to differ in their dispersal time compared to other studied species (Xue et al. 2009). In the lab, all larvae stopped swimming and were exploring the benthos by 19 hours after release, and all had settled by 43 hours. In unfavorable conditions the larvae may travel farther: under high artificial illumination (which increased mortality), sponge larvae swam for a maximum of 24 hours and some were still exploring the benthos when the experiment was terminated at 68 hours. These data are consistent with the larval ecology of other sponges (Maldonado 2006). It therefore seems unlikely that the larvae of this species have exceptionally higher dispersal than to other sponges.

The data do not seem sufficient to form a strong hypothesis about the native range of this species. It seems unlikely to be California, as almost no genetic variation was found at the ITS or cox1 loci in that region. Moreover, the species has likely undergone range expansion in California between the 1920s and the present. Europe is perhaps the most likely source, as we know it was present there in the early 1800s. The genetic diversity at cox1 in Portugal seems notably higher than the diversity at the ITS locus in Asia, though better support would clearly come from comparing data from the same locus.

Future work will be needed to understand what impact this species has on host ecosystems. Its abundance in some habitats seems to make impacts likely, if for no other reason than the occupation of space (Wasson et al. 2001). It is also notable that it has successfully colonized the kelp forests in California, which have been relatively resistant to invasion (Steneck et al. 2002). Within these kelp forests, my observations suggest that this species can be monitored, if imperfectly, using roving diver surveys or photo transects. Though some sponges would require follow-up confirmation in the lab, this might allow the existing large-scale monitoring efforts in California to include this species (Claisse et al. 2018; Miller et al. 2018).

The work presented here builds on the excellent previous work of many authors, and aspects of the pattern I describe have certainly been recognized by others. Hoshino et al. (2008) and Gastaldi et al. (2018) both remarked upon the similarity of *Hymeniacidon* in the *H. perlevis* clade, though in both cases they referred to this clade as a species complex that might be synonymized in the future. Others have simply started referring to their samples by the senior synonym *H. perlevis*, even if they are within the range ascribed to one of the other taxa (Xue et al. 2009; Gastaldi et al. 2018). I build on these earlier efforts by adding data from Southern California and, for the first time, presenting all the genetic data from other projects in one analysis. I recommend synonymizing *H. sinapium* and *H. perlevis*, and recognizing that *H. perlevis* is an invasive species with a global distribution. My hope is that recognition of the unusual distribution and abundance of this species motivates further work into its ecology and ecological impacts.

## Acknowledgements

I am grateful for the help and support of many people in UCSB’s Marine Science Institute and Diving & Boating Program, especially Robert Miller, Clint Nelson, Christoph Pierre, Frankie Puetzer, Christian Orsini, H. Mark Page, and Alecia Dezzani. Steve Lonhart (NOAA) and Shannon Myers (UCSC) were instrumental in facilitating collections in Central California, and the Natural History Museum of Los Angeles’ DISCO program facilitated collections in Los Angeles County.

